# Selective targeting of mutant *huntingtin* intron-1 improves rescue provided by antisense oligonucleotides

**DOI:** 10.1101/2025.07.29.665998

**Authors:** Robert M. Bragg, Christian Landles, Edward J. Smith, Georgina F. Osborne, Jeffrey P Cantle, Gillian P. Bates, Jeffrey B. Carroll

**Author notes:** **One Sentence Summary:** Selective lowering of *mHtt* and *Htt1a* is superior to non-selective lowering of *Htt*, in rescuing aggregate formation and transcriptional dysregulation.

## Abstract

Huntington’s disease (HD) arises from the toxic gain of function caused by a CAG expansion in the coding region of the *HTT* gene. HD is increasingly appreciated to emerge from multiple pathogenic processes, including somatic instability in mutant *HTT’s* (*mHTT*) CAG repeat tract, which leads to diverse deleterious consequences. These include the alternative processing of *HTT* pre-mRNA to generate the *HTT1a* transcript that encodes the very toxic, mHTT isoform referred to as HTT1a. We set out to compare the efficacy and safety of allele-selective lowering of mHTT compared to non-allele-selective lowering using antisense oligonucleotides (ASOs) in heterozygous *Htt^Q111^* (Q111) mice. We developed a mutant specific ASO (MutASO) targeting *Htt* intron-1 that selectively reduced mutant full-length HTT, as well as HTT1a, in the brains of Q111 mice. Compared to the rescue provided by a pan-allele-targeting ASO (PanASO) that lowers wild-type HTT and full-length mHTT (sparing HTT1a), the MutASO essentially eliminated aggregate formation, and provided marked protection from transcriptional dysregulation in HD knock-in mice. Thus, by targeting the ASO to the region upstream of the cryptic polyadenylation sites required to generate the *HTT1a* transcript, our allele-selective MutASO potently reduced HTT1a protein levels. Here, our findings advocate that HTT1a may have a disproportionate impact on aggregate formation and transcriptional dysregulation and that lowering the levels of HTT1a could provide benefit when designing HTT-lowering based therapeutic strategies for HD.

## INTRODUCTION

Huntington’s disease (HD) is an autosomal dominant neurodegenerative disorder caused by the expansion of a glutamine-coding CAG repeat in the huntingtin gene (*HTT*) beyond a critical threshold (*1*). Onset of HD is generally in mid-life, with a progressive series of cognitive, affective, and movement symptoms with age of onset closely linked to CAG repeat size (*2*).

Genetic evidence implies that HD arises from a toxic gain-of-function, though which perturbation(s) of HTT’s function drive toxicity, or whether the toxic gain-of-function is entirely neomorphic remains unclear (*3*). These derangements in cellular homeostasis are preferentially toxic to neurons, with a marked loss of medium spiny neurons (MSNs) in the striatum, though the origin of this cellular specificity is not completely understood. Given these gaps in our knowledge, it remains a challenge to parse which molecular and physiological changes in HD neurons are on causal pathogenic cascade and which are epiphenomenon.

Because of the imperfect understanding of HD’s pathogenic cascade, lowering levels of mutant HTT (mHTT) protein and/or mRNA (*mHTT*) is an attractive therapeutic approach. The number and chemical diversity of these “huntingtin lowering” approaches have exploded over the previous few years, with many treatments in various stages of clinical development (*4–6*). Current HTT lowering approaches in the clinic include adeno-associated virus (AAV) delivered micro RNAs (miRNA) (*7*), antisense oligonucleotides (ASOs) (*8, 9*), and small molecule splice modulators that reduce *HTT* mRNA via nonsense mediated decay (*10, 11*). A consistent concern about non-selective HTT lowering is the safety of adult-onset reductions in wild-type HTT levels (wtHTT); HTT is highly conserved, widely expressed and has poorly delineated molecular functions (*5*). Strong human genetic evidence suggests that 50% loss of HTT is not associated with any phenotypes in humans (*12–14*), arguing in favor of either a graded or an allele-selective approach to HTT lowering.

During aging, *mHTT*’s CAG repeats are susceptible to expansion-biased somatic instability (SI). This has been clear for decades, in both HD patient tissues and preclinical HD mouse model data (*15*). Recent genome wide association studies (GWAS) have implicated variants in a surprising number of DNA mismatch repair genes in the timing of onset of HD symptoms, as well as *HTT*’s CAG repeat purity (*16–19*). Amongst the cell types examined that are susceptible to SI, MSNs are the most likely and earliest to expand (*20, 21*), though the reasons remain obscure. Expression levels of mismatch repair genes have recently been implicated based on cell-type-specific RNAseq from human post-mortem tissue (*22, 23*). Further, small pool PCR (*21, 24*) and single cell (*25*) analyses suggest that individual MSNs may have dramatic expansions of hundreds of CAG repeats, even in a patient inheriting a CAG size in the low 40s.

Very long CAG repeats in *mHTT* arising from SI could lead to many deleterious effects within the cell. One that is relatively well described is the alternative processing of *HTT* pre-mRNA, in which cryptic polyadenylation (polyA) sequences in intron-1 of *HTT* are activated, leading to the formation of a translationally competent exon-1 only species referred to as *HTT1a* (*26–29*). The generation of this fragment is correlated with CAG length, and age-related SI could lead to individual neurons with very long CAG repeats, and thereby susceptible to increased production of the HTT1a protein within that cell (*29, 30*). The HTT protein is primarily cytoplasmic (*31*), and yet nuclear aggregates containing mHTT have long been observed, particularly in striatal neurons in HD mouse models (*32*) and patients (*33*). Undoubtedly HTT (or N-terminal fragments of it) must enter the nucleus to form aggregates, and one proposal for HTT1a’s high toxicity is the known ability of this HTT isoform to translocate into the nucleus, preferentially seed aggregate formation, and drive transcriptional dysregulation (*34, 35*).

We have a long-standing interest in the most parsimonious means of conducting HTT lowering using ASOs to preferentially silence *mHTT* while sparing wild-type *HTT (wtHTT)* (*36–38*). Here, we developed a set of ASOs to selectively lower *mHtt* (MutASO) or target both *Htt* alleles (PanASO) in the context of heterozygous *Htt^Q111^*HD model mouse with ∼111 CAGs (hereafter referred to as Q111) (*39*). Treatment with both ASOs leads to potent and long-lasting reductions in the protein(s) arising from the targeted allele(s). Fortuitously, our MutASO targets a region in intron-1 of m*Htt* upstream of the cryptic polyadenylation sequences needed to produce HTT1a, and robustly reduced levels of the HTT1a mRNA and protein. This was correlated with a dramatically improved rescue of HD-relevant pathological signs compared to parallel PanASO treatment, notably nuclear aggregates of mHTT and improvements in transcriptional dysregulation. Our results suggest that allele-selective targeting of *mHtt*’s intron-1 to decrease levels of HTT1a as well as full-length mHTT has surprisingly large benefits for HD compared to full-length pan-lowering of both wtHTT and mHTT.

## RESULTS

### Identification and Characterization of Allele Selective ASOs for Q111 Mice

The *mHtt* allele in the Q111 mouse was created by replacing a 296-bp mouse exon1-intron1 fragment with a 590-bp human exon1-intron1 fragment via homologous recombination (*39*). This created a chimeric human-mouse *Htt* exon-1 containing an expanded CAG, as well as 268-bp of humanized sequence in intron 1 of the Q111 allele compared to wt*Htt* allele (Fig. 1A). We used this sequence to design a set of ASOs targeting *mHtt* transcripts while sparing endogenous *wtHtt*. To identify ASO sequences, we designed 22 ASOs targeting the human-specific knock-in sequence in this region. All molecules tested had asymmetric 2′-ribose modifications and rational-designed stereopure phosphorothioate (PS) backbone chemistries previously demonstrated to enhance the potency of antisense oligonucleotide (*40, 41*). Initial sequences were screened in human iCell neuronal cells (in which both alleles are targetable, as the target sequence is human) and potency for human *HTT* knockdown was ranked (Fig. 1B). The most potent ASOs were then screened in primary neurons from Q111 mice, with HTT protein levels and knockdown measured by western blot to distinguish wtHTT versus mHTT species (sample data in Fig. 1C). These blots reveal robust and selective knockdown of mHTT and relative sparing of wtHTT. We then generated MutASO-1101 (hereafter MutASO) by introducing phosphoryl guanidine (PN) backbone linkages (*42*) to the top 2 ASOs, MutASO-4 and MutASO-11 and tested *in vivo* (Fig. 1D). MutASO resulted in 68.3% mHTT KD at 4-weeks, as measured by an antibody-based electro-chemiluminescence Mesoscale Discovery (MSD) assay (*43, 44*) and resulted in no induction of neurofilament light chain (NEFL) levels in the plasma (Fig. 1E-F). We then modified a published mouse/human cross-reactive ASO targeting exon-42 (*45*) with the same 2’-ribose chemistry as our MutASO and rationally designed PS-backbone stereochemistry resulting in a “PanASO” that targets both alleles of *Htt* in the Q111 mouse (Fig. 1D). Notably, the region of humanized knock-in sequence in our mice obligatorily places the MutASO target sequence upstream of cryptic polyA sites in intron-1 that enable generation of the HTT1a splice variant of *mHTT* (*26*).

**Figure 1.**
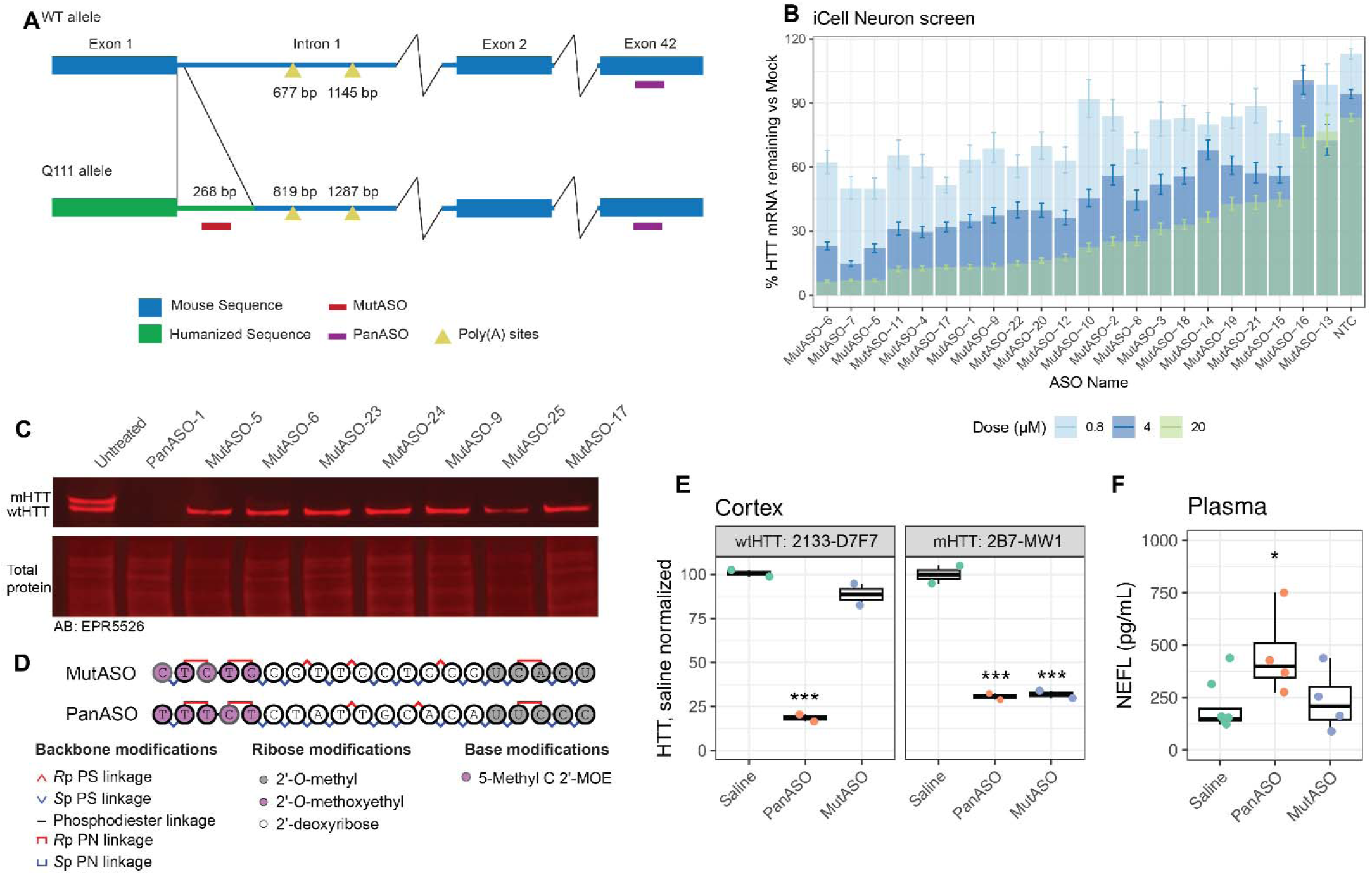
Development of mutant allele-selective ASOs in the context of Q111 mice. (**A**) A schematic depicting the humanized mHtt sequence of the Q111 allele compared to the wtHtt mouse allele, relative to the described cryptic poly-adenylation sites in intron-1. The mutant-and pan-selective ASO targets are indicated. (**B**) Htt mRNA levels in iCell Neuron after 7 days of treatment with 0.4 (light blue), 4 (blue) or 20uM (green) ASO targeting human intron 1 sequences (ASO n=2 replicates, NTC n = 8, with 2 technical replicates for all samples). (**C**) Primary neurons from Q111 mice were measured for HTT by western blotting (EPR5526) after 7 days of treatment with the indicated ASO. (**D**) Final ASO nucleotide sequences and backbone modifications including chiral right-(Rp) and left-(Sp) handed linkages, phosphorothioate (PS), phosphodiester, and phosphoryl guanidine (PN) backbone chemistry (**E**) Protein levels of wtHTT and mHTT with the indicated MSD assays in the cortex of Q111 mice treated with 300 µg of ASO via unilateral ICV injection (N=2/group). (**F**) Plasma NEFL levels were established with an MSD assay 28 days after ICV injection of the indicated compound (N=4-6/group). Data in B are mean +/-SEM and data in E, F are boxplots with individual points representing each mouse and were analyzed by factorial ANOVA with Tukey post-hoc test. ***P< 0.01, *P<0.05, compared to saline.

### Interim Silencing Results and Overview of Interventional Study Design

Based on our previous power analysis of endpoints in Q111 mice, we initiated an interventional study and treated mice from 2.5 -8.5 months of age, a time over which several translationally relevant phenotypes emerge in this model (*46*). Washout studies with both ASOs administered by unilateral intracerebral ventricular (ICV) injections were conducted to determine the dosing interval over which the reduction in HTT levels could be maintained to minimize re-administration of the treatment (Fig. 2A). These results led to our re-treating mice with ASOs 3-months (13 weeks) after initial treatment and sacrificing them 3-months (13 weeks) following the second injection; giving a total treatment duration of 6 months (Fig. 2B) with the intention of maintaining HTT level reduction of 50-75% for the full study duration.

**Figure 2.**
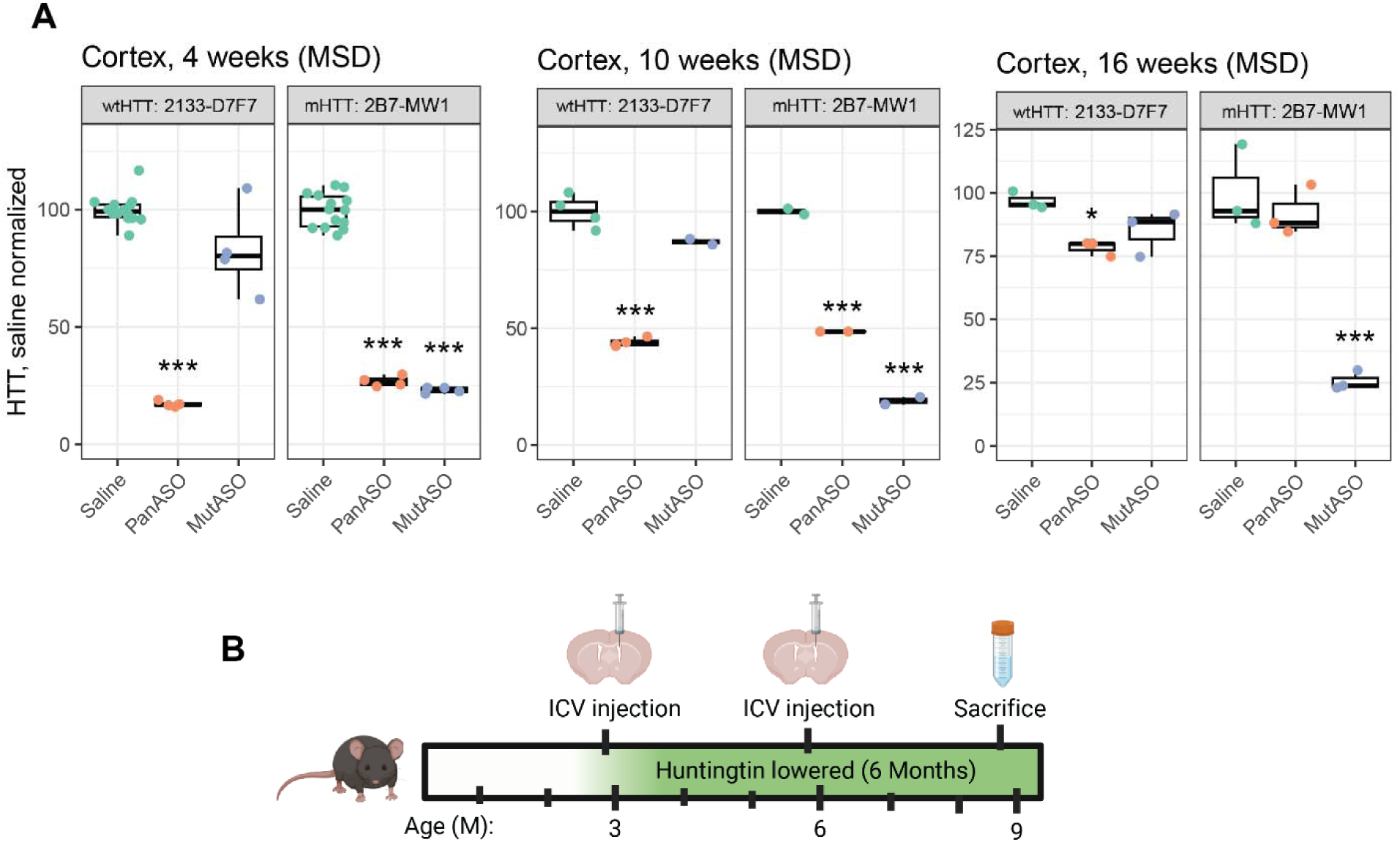
Overview of interventional study design based on the duration of action of each ASO. (**A**) Levels of wtHTT and mHTT in the cortex of Q111 mice at 4-(left), 10-(middle) and 16-weeks (right) after a single unilateral ICV injection of indicated treatment (N=3-15). (**B**) A schematic overview of the trial design indicating the timing of injections and sacrifice. Data in A are shown as boxplots with individual mice plotted as points and were analyzed by factorial ANOVA with Tukey post-hoc test. *P < 0.05 and ***P < 0.001 for comparison to saline.

As a measure of neurotoxicity, we monitored NEFL levels both in pilot studies and in the complete longitudinal study. We observed a relatively mild 2.3-fold induction of NEFL at 4-weeks by PanASO (Fig 1F). However, we did not observe any increase after the full study was complete. In contrast, we observed a 3.5-fold NEFL induction of the MutASO in both WT and Q111 mice (Fig. S1). Whilst Q111 mice do not generally exhibit any robust behavioral changes until 14-18 months of age (*47*), we assessed their overall activity levels using an open field behavioral assessment, to determine whether treatment with either ASO lead to hypo-or hyper-activity. While overall ANOVA for both genotype and treatment were significant (p<0.05) for genotype and treatment, post-hoc analyses no differences between genotypes at any treatment, including saline at 8.5 months of age (Fig. S2A-B).

### HTT levels following PanASO and MutASO treatment were reduced

Mice were sacrificed at 8.5 months of age (6 months post-treatment) and tissues were collected for analysis. We quantified HTT levels using MSD across the whole cohort. As expected, PanASO reduced both wtHTT and mHTT in the cortex of WT and Q111 mice (Fig. 3A). In WT mice, wtHTT was reduced by 30% and in Q111 mice, wtHTT was reduced by 43% and mHTT was reduced by 45%. Treatment with MutASO reduced mHTT by 86% in Q111 mice, while sparing wtHTT in both WT and Q111 mice (Fig. 3A). To establish the distribution of knockdown more broadly, we measured full-length mHTT in a wider set of brain tissues using another sensitive antibody-pair based method: homogenous time-resolved fluorescence (HTRF; Table S1) (*30, 48*). Using an assay specific to full-length mutant HTT, we confirmed an approximately 50% knockdown of mHTT in the PanASO groups (Fig. 3B; striatum: 43.3% reduction; cortex: 44.6% reduction; cerebellum: 50.3% reduction) and robust reduction of mHTT in the MutASO treated groups (Fig. 3B; striatum: 91.3% reduction; cortex: 92.5% reduction; cerebellum: 71.5% reduction). The general observation of reduced mHTT protein detected by HTRF was visually confirmed in cortical lysates by immunoprecipitating with an antibody from targeting expanded polyQ protein and immunodetecting with an antibody targeting an epitope at around 1,218 amino acids (Fig. 3C) (*49*).

**Figure 3.**
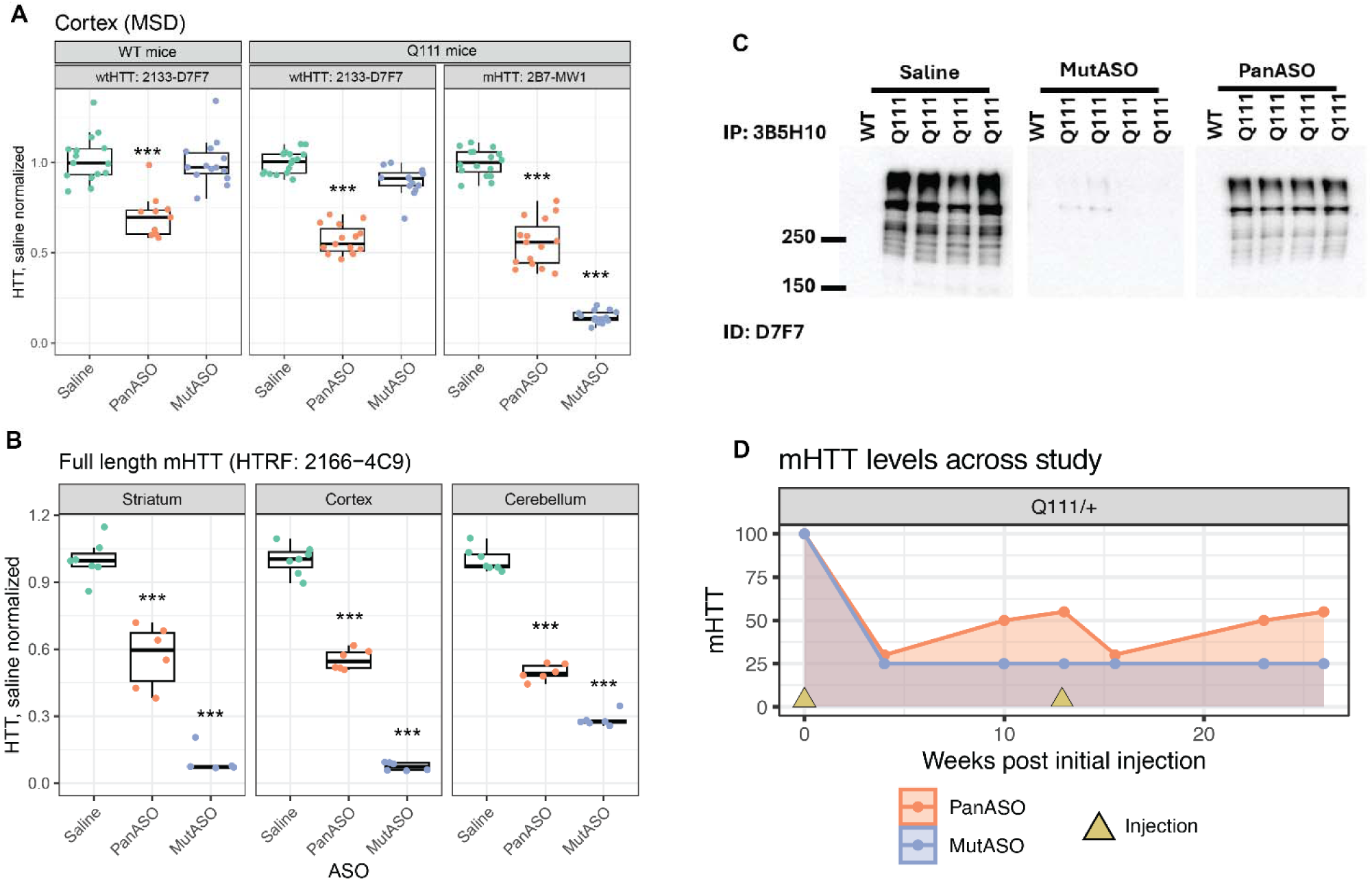
Both wtHTT and mHTT levels were reduced at the end of our interventional study. (**A**). HTT levels measured by MSD in our entire interventional cohort measured at 26 weeks show significant reduction of wtHTT and mHTT in response to PanASO. Additionally, MSD shows no reduction of wtHTT in response to MutASO (WT mice: p = 0.99, Q111 mice: p = 0.06), however there was a strong reduction of mHTT in the Q111 mice in response to MutASO (n = 11-16 / group). Specific antibody pairs are indicated in each panel (**B**). Additional analysis of mHTT in a wider set of brain regions measured by HTRF confirms knockdown of mHTT in Q111 mice in response to both PanASO and MutASO in the striatum, cortex and cerebellum at 26 weeks (n = 7 for saline, 6 for ASO groups). (**C**). Reductions in cortical mHTT measured by HTRF are confirmed by immunoprecipitation (IP) and immunodetection (ID) with indicated antibodies (n = 04) (**D**) Approximate levels of mHTT knockdown across the study based on separate experimental data from both Fig 2A (shown as data points at 4 and 10 weeks following each injection) and endpoint levels at week 26 from B. Data in A and B are presented as boxplots with individual mice as points and were analyzed by factorial ANOVA with Tukey HSD post-hoc test. ***P < 0.001 for comparison to saline.

Importantly, by design, these measurements capture HTT levels at the furthest time from injection possible (13 weeks post-injection). Based on our washout studies (Fig. 2A), we infer that mHTT levels were equally suppressed from treatment until at least 4 weeks of age, and that the PanASO treatment resulted in a slowly increasing amount of both HTT isoforms from 5-13 weeks. The mice were then re-injected with ASO and experienced a similar trough and slow rebound of HTT. In contrast, the MutASO remained nearly equipotent from 0-16 weeks of age – a graphic of our model of mHTT levels is illustrated in Fig. 3D.

### MutASO, but not PanASO, Reduces HTT1a Formation in Q111 Mice

As CAG tracts expand, they become sensitive to the generation of an additional translationally competent exon-1 only transcript (“*Htt1a*”, Fig. 4A). The HTT1a protein enters the nucleus, is notably toxic compared to other N-terminal fragments and avidly seeds aggregates (*28*). We measured both soluble and aggregated forms of HTT1a using HTRF assays (*48*). We found that the striatum, cortex, and cerebellum had similar patterns of soluble HTT1a expression (Fig. 4B). In general, we detected an increase of soluble HTT1a following PanASO treatment compared to saline treatment (6-39%). This increase was somewhat surprising, given the reductions in full-length mHTT, however our current understanding is that a reduction in full-length mHTT allows for better detection of the soluble HTT1a fragment (Papadopoulou *et al*., this issue). In contrast, we detected lower soluble HTT1a levels following MutASO treatment compared to saline treated mice across all regions tested (Fig. 4B). Using an antibody pair selective for aggregated HTT1a (*28, 50*), we found no significant reduction of aggregates in the striatum of PanASO treated mice (Fig. 4C). In contrast, the aggregated striatal HTT1a signal was reduced by 68% and was comparable to HTT1a signal from 2-month-old ‘baseline’ mice, that have negligible levels of aggregated HTT1a (*30*). While most aggregation in the brains of Q111 mice at 8.5 months of age was in the striatum (Fig. 4C), we did see a modest reduction of aggregated HTT1a from PanASO treated mice in the cortex (28%), however, there were much stronger reductions from the MutASO treated mice in both the cortex and cerebellum (Cortex: 58%; Cerebellum: 26%; Fig. 4B).

**Figure 4.**
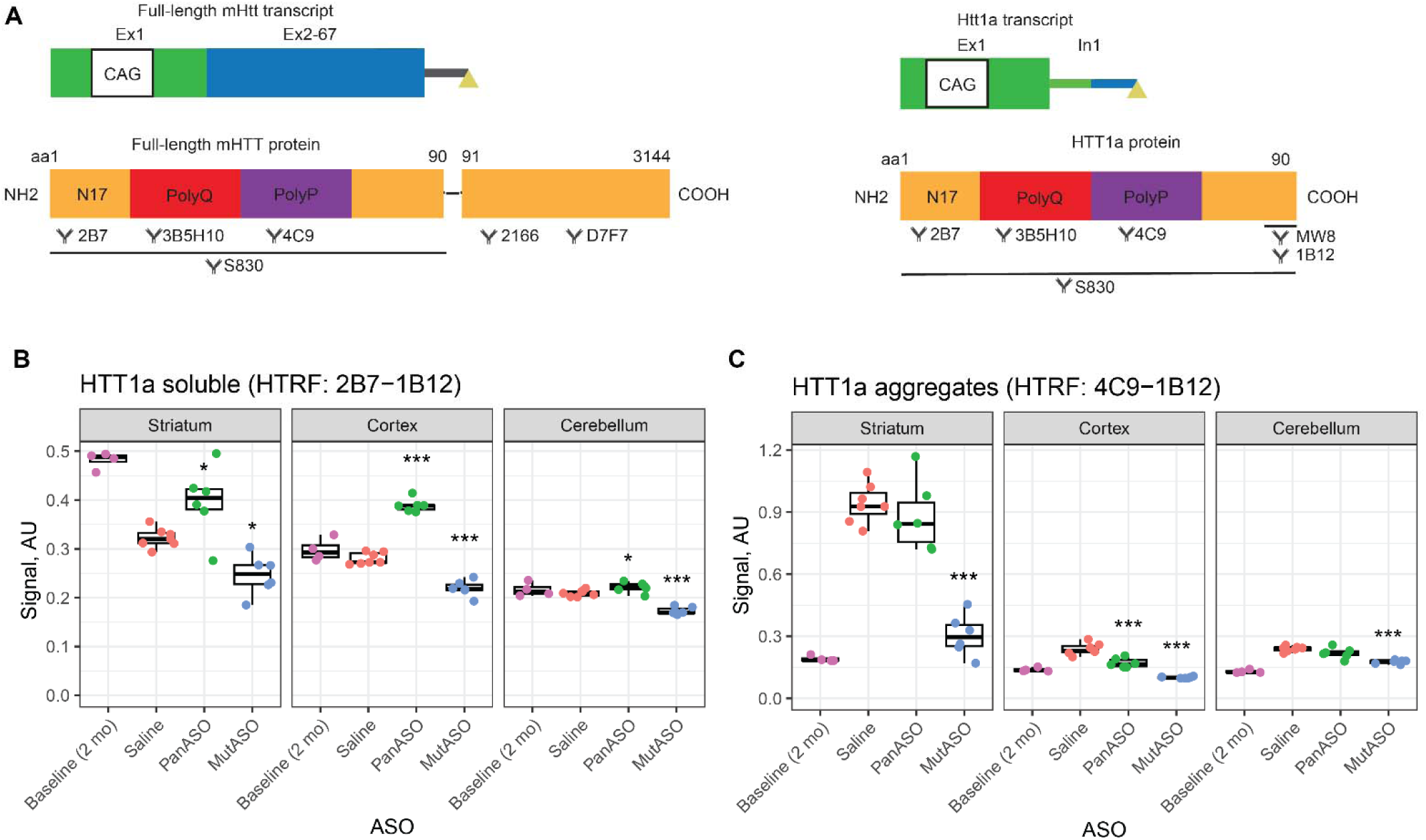
MutASO, but not PanASO, reduces soluble and aggregated HTT1a fragments. (**A**). Schematic of full-length HTT and HTT1a’s transcript and the epitopes of protein structure and antibodies used throughout. (**B**). Soluble levels of HTT1a were detected by an HTRF assay using the 2B7 and 1B12 antibody pair revealing soluble HTT1a levels were reduced by MutASO treatment, but not by PanASO, in the striatum, cortex, and cerebellum. (**C**) Aggregated HTT1a was detected by an HTRF assay using the 4C9 and 1B12 antibody pair. At sacrifice, aggregated HTT1a was reduced in the striatum, cortex and cerebellum by MutASO. Samples from untreated mice serving as a baseline level at the approximate age of ASO treatment onset (2-month-old) are shown to indicate the expected amount of each species across tissues at study initiation but are excluded from statistical analyses. Data are presented as boxplots with individual mice as points and were analyzed by factorial ANOVA with Tukey HSD post hoc-test. *P<0.05, ***P < 0.001 for comparison to saline.

### MutASO, but not PanASO, Rescues HTT Aggregate Formation in Q111 Mice

To confirm the reductions in aggregated HTT species measured by HTRF, we completed immunostaining in the striatum of Q111 mice. MW8 is a conformation-specific antibody that is specific for HTT1a under certain conditions (*49*). By immunohistochemistry, it detects inclusion bodies in neuronal nuclei, but not diffuse aggregation (*46, 51–53*). Using MW8, we observed the expected distribution of large HTT inclusions in the striatum of saline treated Q111 mice (Fig. 5A). Remarkably, treatment with PanASO did not reduce the abundance of inclusions, compared to saline treated mice – in both cases large inclusions were present in approximately 25% of striatal neurons. In contrast, MutASO mice showed a near-complete loss of large HTT inclusions, where they were present in only 1% of neurons. We next used the S830 antibody that in addition to detecting HTT1a aggregates, also detects aggregates containing N-terminal proteolytic fragments of mHTT, should they be present. The S830 antibody also detects the diffusely aggregated HTT, which is the first aggregated species to form in neuronal nuclei and that coalesces into inclusion bodies with disease progression (*34, 54*). We quantified the area of neuronal nuclei positive for S830 reactivity, finding that it covers an average of 11% of each neuronal nuclei in the striatum of saline and PanASO treated mice. In contrast, S830 positive HTT immunoreactivity covers less than 1% of each striatal nuclei in MutASO treated mice. These IHC results suggest that both diffuse HTT aggregation and large nuclear inclusions are eliminated from MutASO treated mice, consistent with the proposed model that it is HTT1a that nucleates HTT aggregation in the nucleus (*54*).

**Figure 5.**
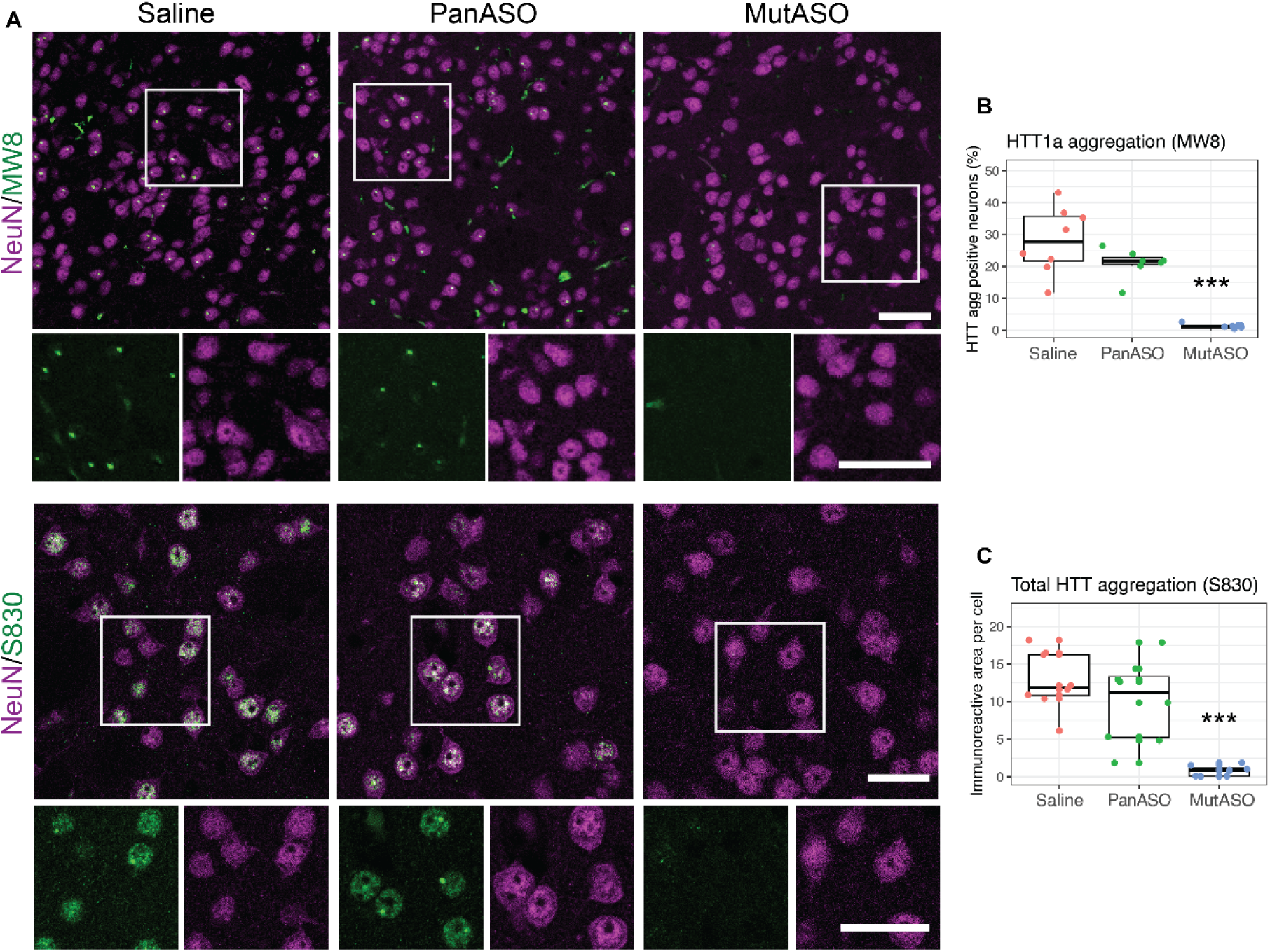
Immunohistochemistry for mHTT neuronal nuclear aggregates in the striatum reveals a striking effect of MutASO, but not PanASO, treatment. (**A**) IHC from mice indicated treatment using a HTT1a-specific aggregate (MW8; scale bar = 75µm) and a total HTT aggregate (S830; scale bar = 30µm) show marked reduction in immunoreactivity in MutASO groups but no obvious change in PanASO. (**B**) Quantification of MW8 (n = 8/group) and (**C**) S380 (n = 8, 2 images per group) reveal a near complete loss of aggregate positive neurons (B) or area per cell positive for immunoreactivity (C). Data are presented as boxplots with individual mice as points and were analyzed by ANOVA with Tukey HSD post hoc-test. ***P < 0.001 for comparison to saline.

### MutASO, but not PanASO, Rescues Transcriptional Dysregulation in Q111 Mice

It has been proposed that HTT1a-seeded aggregate formation is a critical, rate-limiting, step in transcriptional dysregulation, and thereby HD pathogenesis (*34*). To test this hypothesis, we conducted bulk RNA sequencing from the striatum. In the saline-treated Q111 mice, we observed a very close recapitulation of the transcriptional dysregulation that we (*46, 55*) and others (*56*) have repeatedly observed in aging Q111 mice, helping to define a core signature of HD-relevant differential expressed genes (DEGs; Fig. 6A) (*57*). Universally, if total HTT reduction is effective in ameliorating transcriptional dysregulation in HD model mice, we would expect to see restoration of Q111 striatal gene expression signatures towards WT mouse levels in PanASO-treated mice. Instead, expression of only two genes was altered by PanASO treatment, one of which being *Htt* itself (Fig. 6B). This was in sharp contrast to the MutASO treatment, which led to a very marked shift in the striatal transcriptome of Q111 mice (Fig. 6C).

**Figure 6.**
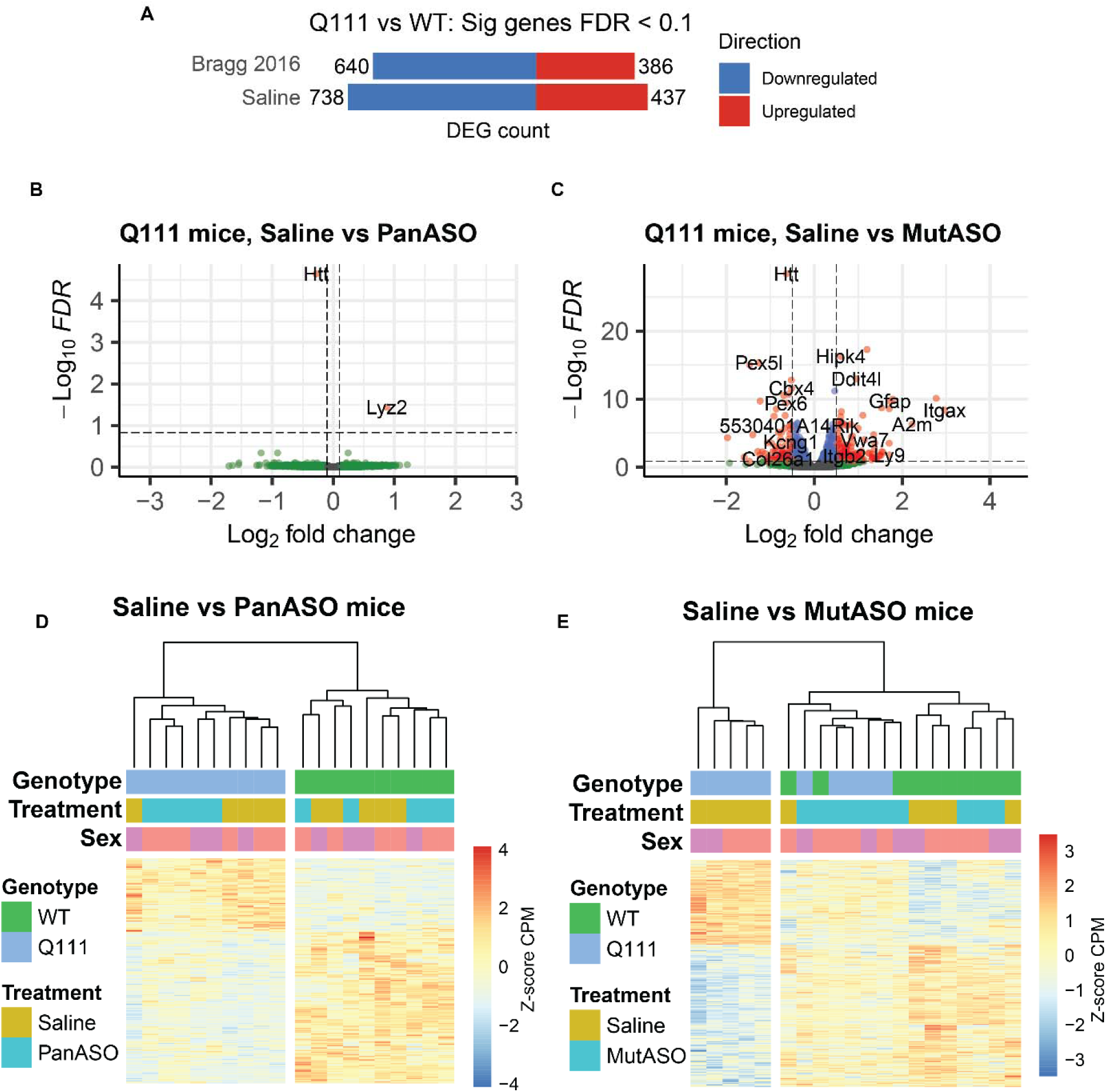
MutASO, but not PanASO, rescues broad patterns of transcriptional dysregulation in the striatum of Q111 mice. (**A**) The number of up-and down-regulated striatal DEGs in this study was compared with historical data from similar aged mice (9-months) in Bragg et al., 2016. **B, C**) Volcano plots indicating the changes induced by PanASO in the Q111 mice used in this study – only two genes are altered and MutASO in Q111 mice where 315 genes are significantly altered. (**D, E**) Heatmaps of gene expression produced by unsupervised hierarchical clustering of the 1175 Q111 induced DEGs. Gene expression is shown as Z-score normalized counts-per-million.

To assess whether these DEGs reflect a normalization of gene expression, rather than an exacerbation, we turned to unsupervised clustering of the ∼1,200 DEGs between treatment groups (Fig. 6D-E). This approach attempts to cluster mice (columns) to reduce the variance in the total set of differentially expressed genes (rows). Post-hoc labeling of the categories of interest (genotype and treatment, in this case) enables a gestalt assessment of global changes.

Focusing on comparing the impact of saline treatment to that of PanASO, genotypes of mice are very neatly split, with no clear impact or clustering driven by the treatment group, indicating no large-scale restoration of gene expression after PanASO treatment (Fig. 6D). In contrast, every individual MutASO-treated Q111 mouse clustered with WT mice, suggesting a large shift towards normalization of both up-and down-regulated DEGs (Fig. 6E). (*58*)

Following a recently developed more manual approach, we classified 188 previously defined core HD signature genes by the extent of their rescue or exacerbation from ASO treatment (*57, 58*). Treatment with MutASO led to 100 of 188 HD signature genes being partially or completely rescued, whereas treatment with PanASO led to zero rescued genes (Fig. 7A). Manual examination of several critical striatal-specific core HD DEGs reveals remarkable protection from MutASO, but not PanASO, treatment (e.g. *Ppp1r1b*, *Penk*, *Scn4b*, etc.; Fig. 7B). Finally, we compared the enrichment terms from saline treated Q111 vs WT DEGs (i.e., a baseline GO enrichment signature) to the signature from MutASO treated Q111 vs Saline WT DEGs (Fig.7C). As expected, we see that the baseline enrichment terms are dominated by neuronal function (e.g., regulation of synapse organization, transporter activity) and see that the p-values and gene count enrichments are lower for all categories in the MutASO treated mice, highlighting a coherent rescue of HD signature related pathways.

**Figure 7.**
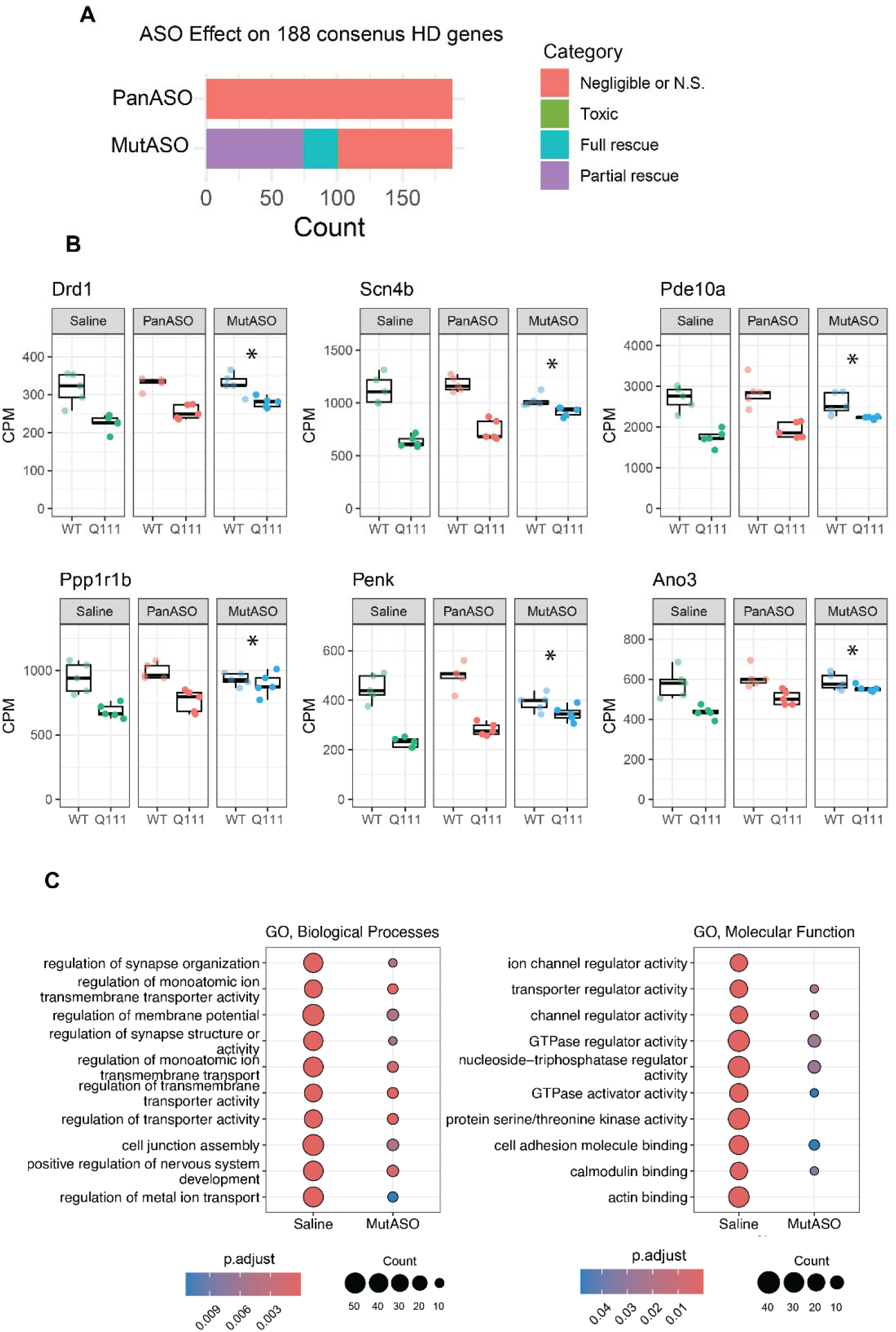
Treatment with MutASO, but not PanASO, robustly protects from core HD DEG dysregulation. (**A)** 188 genes comprising a common HD signature were categorized into 4 bins. To be considered rescued by ASO, DEGs from the Q111+Saline vs Q111+ASO comparison (FDR < 0.1) that were also significant from the Genotype x Treatment interaction (P<0.05) were binned into categories based on expression compared to the saline control. No genes met these criteria for the PanASO treatment. For MutASO, 26 genes were fully rescued (expression delta between Q111 Saline vs Q111 ASO was 85-115% of Saline WT value), 74 genes were classified as partially rescued (expression delta between Q111 Saline vs Q111 ASO was within 15-85%f Saline WT expression), 1 gene was exacerbated (expression delta was >115% of saline WT), and 87 genes showed no rescue (FDR > 0.1 or expression within 15% of saline WT). (**B**) A small panel of genes from A. * indicates partial rescue based on the statistics outlined in A. Data are presented as boxplots with individual mice shown as points. (**C**) Gene ontology (GO) pathway enrichment of DEGs (FDR < 0.1) from saline treated Q111 vs WT mice and MutASO treated Q111 vs Saline WT mice. The top ten GO biological processes (left) and GO molecular function (right) pathways from the baseline HD signature are shown with the corresponding enrichment following MutASO treatment. Reductions in the number of genes and enrichment p-value following treatment are significant for all pathways shown (z-test adjusted p < 0.01).

To assess the potential off target impact of MutASO on transcription, we examined genes that were altered by MutASO in WT mice. We found a small signature of 119 genes, 84 of which were well annotated (Fig. S2A, FDR < 0.1 and absolute log2FC > 0.5). When assessed for over enriched KEGG pathways using EnrichR, we find only one significantly enriched pathway, ‘complement and coagulation cascade’ with 5 of 88 genes overlapping (*P_adj_* = 0.02, Fig. S2B), suggesting this ASO induces a relatively minor immune response. The PanASO only induced one altered gene beyond *Htt* (Fig. 6B), *Lyz2*. This gene was among the off-target MutASO genes, suggesting that its differential expression was likely a response to the ASO itself, rather than *Htt* lowering.

## DISCUSSION

In this study we compared the rescue obtained by treating HD mice with allele-selective ASOs targeting mHtt’s intron-1 to non-allele-selective ASOs targeting exon-42. We observe striking benefits provided by our allele-selective intron-1 targeting ASO, including a reduction in HTT1a levels and a marked phenotypic improvement compared to targeting downstream. Treatment with an ASO targeting intron-1 led to nearly complete abrogation of HTT aggregates, and a robust rescue of transcriptional dysregulation. To our knowledge, this study is the first to report robust transcriptional rescue in HD mice after ASO treatment.

Notably, we find that treatment with MutASO led to marked reductions in both the soluble and aggregated form of the HTT1a isoform (Fig. 4). The initial observations supporting the existence of the *HTT1a* transcript were made more than 10 years ago (*26*), and interest in HTT1a has increased over time, as it is known to be highly prone to aggregation (*59*) and toxic (*60*). The production of *HTT1a* correlates to CAG repeat length, such that longer CAG repeats drive the more frequent activation of cryptic polyA sites and the alternative processing of the *HTT* pre-mRNA (*29*). There are at least two non-exclusive interpretations of the markedly improved rescue with MutASO -first, that allele-selective mHTT lowering is markedly superior to pan-HTT lowering due to maintained expression of wtHTT. Second, MutASO’s ability to reduce HTT1a provides protection out of proportion to its ability to reduce full-length mHTT levels. We believe our data are more consistent with the latter model, given the phasic shift in improvement in rescue of HTT aggregation (Fig. 5) and transcriptional dysregulation (Figs. 6-7) – the difference in rescue between the ASOs is not graded by the extent of HTT knockdown, but rather a step function, in terms of the size of the rescue after HTT1a knockdown. This abruptness argues in favor of distinct molecular mechanisms underlying these rescue effects, though not definitively. Unfortunately, using gapmer ASO designs as we have here, it is not possible target HTT1a selectively without concomitantly reducing full length mHTT, so additional data are required to definitively determine the role of HTT1a in the rescue we observe in our study.

The simplest interpretation of the fact that many MMR genes are accelerating modifiers of HD onset (*16, 17, 61*) is that their collective action drives enhanced SI in *HTT*’s CAG repeats. Recent findings have lent complexity to this simple model – some generally supportive of it, such as single-cell profiling of both CAG sizes and transcriptional profiles in HD patient striatal neurons (*25*); these reveal a striking array of CAG sizes in individual MSNs, with tight predictability of transcriptional changes by CAG lengths. This supports older, bulk, assessments of tissue from both patients with HD and mouse models (*62, 63*). In contrast, after sorting specific cell types from the brain of patients with HD, no simple relationship was found in SI profiles in pseudobulk analysis of pooled cells of a given type, and some relatively preserved cell types were shown to experience somatic instability, and vice versa (*64*). Further complexity is added by the recent expansion of the GEM-HD consortium’s expanded GWAS with nearly 12,000 subjects, which unexpectedly revealed complex relationships between MMR variants, SI and clinical features of HD (*61*). Notably, while variants in some MMR genes (e.g. *MSH3*) modified both SI and HD risk in a coherent way, other MMR gene variants exhibited a much more complex, putatively cell-type-specific, impact on SI. Importantly, despite the huge numbers of subjects under investigation, not a single hit has been observed that clearly modifies HTT/*HTT* levels, and the authors note: “*…, the possibility that full-length mutant huntingtin is not the cause of neuronal dysfunction and death must be considered, so the pathogenic role, if any, of a protein product produced from the expanded HTT allele is of particular interest.*” Our results provide supportive evidence in favor of the potentially preferential toxicity of the HTT1a protein isoform.

There are important limitations to our study. Notably, because ASOs work in both the nucleus and cytoplasm, we were unable to selectively target pre-versus mature mRNA, preventing differentially targeting of HTT1a (*65*). At the onset of this study, our intention was to conduct a simple side-by-side comparison assessing the impact of allele-selective silencing versus pan-HTT lowering. Two factors emerged during the study that challenged our ability to make those comparisons as robustly as we had hoped. First, we discovered that treatment with MutASO knocks down HTT1a, as well as full length mHTT (Fig. 4). Second, we observed a small difference in the half-life of the Pan-and MutASOs which resulted in a period of several weeks in which mHTT was reduced more in the MutASO compared to PanASO (Fig. 3D). We think that the total duration of this discrepancy in knockdown was limited to several weeks, and partial, and therefore unlikely to explain our results. Fortunately, unbeknownst to us, our colleagues were working on a publication with complementary strengths and limitations in parallel (Papadopoulou *et al.*, this issue). They demonstrated that siRNA-mediated lowering of HTT1a in the zQ175DN model of HD provides robust rescue of HTT aggregation and the associated HD-associated transcriptional signature, while lowering of full-length HTT provides almost no impact on these phenotypes. In contrast to our current study, which demonstrated persistent HTT1a and mHTT knockdown with our MutASO and less persistent knockdown of HTT by our PanASO, they achieved strong full-length HTT KD (65%) but relatively weaker HTT1a KD (45%). This complementary study bolsters our interpretation that HTT1a targeting provides robust rescue, despite lower knockdown.

An important caveat with our study is that it was specifically designed to be “allele selective” only in the context of the Q111 mouse model of HD, to compare rescue provided by m*Htt*-selective and non-selective ASOs in a consistent, albeit artificial, genetic background. The MutASO used here would not be allele-selective in HD patients, since the intron-1 sequences are equivalent in HD and non-HD *HTT* alleles (*53*). Conceptually, achieving a m*HTT*-selective oligonucleotide therapeutic targeting intron-1 that relies on targeting heterozygous genetic variations such as single nucleotide polymorphisms (SNPs) within the intron-1 region is possible. However, SNPs are only expected every ∼5kb in human autosomes and identifying common heterozygous SNPs within the several thousand base pairs of targetable sequence in HTT1a’s coding sequence is statistically unlikely (*37, 38, 66*). A more promising approach may be to target specific structural changes or splicing sites that give rise to *HTT1a*, as has been developed for C9orf72 (*40*).

An additional limitation to this study is that we do not report any behavioral rescue experiments. A limitation of HD knock-in mouse models is that the ones that effectively model the earliest molecular changes seen in HD, such as the Q111 mice used here, have only modest and very late behavioral changes, with no overt striatal atrophy or cell death (*47, 67*). On the other hand, models such as the zQ175DN mice have more pronounced and earlier HD-relevant behavioral and neuropathological symptoms (*68, 69*), but have recently been shown to be insensitive to rescue by blocking somatic instability (*70*), suggesting they model the post-SI epoch of mHTT toxicity. Therefore, in a single mouse study of less than 14 months, it is very difficult to provide evidence about both early molecular changes (e.g. nuclear specific mHTT aggregation, transcriptional dysregulation) and behavioral phenotypes. We believe the face validity of MSN-specific nuclear mHTT aggregation and transcriptional dysregulation is high, given that both mice and humans experience these phenotypes in an age-dependent and MSN-specific manner (*71*), and so have made the choice to focus our rescue work on these phenotypes, rather than mouse behavioral ones.

## MATERIALS AND METHODS

### Study design

For the main interventional study, 91 mixed sex littermate mice from two genotypes (WT and heterozygous Htt.Q111) were split across 3 treatment groups (Saline, PanASO, MutASO) and injected with ASO at 2.75 months (85 days) and 6 months (175 days). Due to attrition, 86 mice were sacrificed at 8.5 months (265 days), and tissues were collected for analyses. A detailed breakdown is provided in Table S2. Following injection, researchers were blind to treatment groups.

### Mice and ASO administration

We used heterozygous Q111 mice (Jax ID: 003598) on a C57BL6/J background and WT littermates for all of the studies described here(*72*). For pilot and screening studies, we used mice bred at the Western Washington University (WWU) vivarium. For the interventional study, mice were bred at Jax (Bar Harbor, Maine) and shipped to WWU 3-weeks prior to the start of the study. Mice were CAG sized with an average CAG length of 117 ± 3. Mice were fed ad libitum and maintained with a 12/12 light/dark cycle. Unilateral ICV bolus injections, which result in efficient brain-wide distribution of ASO, were conducted as previously described (*73*). Briefly, mice were anesthetized with isoflurane, placed on a stereotaxic device, and unilaterally injected with 300 µg of ASO in 10 µL of PBS at 2.5 and 6 months of age. All animal work was approved by the WWU animal use committee (protocol 22-001).

### ASO synthesis

We synthesized and purified chemically modified, stereopure oligonucleotides as described with minor modifications (*42, 74*). Oligonucleotide purity was characterized by LC–HRMS and HPLC. MutASO sequence is 5’-CTCTGGGTTGCTGGGUCACU-3’ and PanASO sequence is 5’-TTTCTCTATTGCACAUUCCC-3’ with chemistry and modifications indicated in Fig. 1D.

### In vitro ASO screens

iCell GABA Neurons 01434 (FUJIFILM Cellular Dynamics) were plated on Matrigel (Corning) coated 384-well plates using the Bravo liquid handling platform (Agilent) at 40,000 cell/well. Cells were cultured according to manufacturer’s recommendations in iCell Base Medium plus iCell Neural Supplement A (FUJIFILM Cellular Dynamics). 24 hours after plating, the media was replaced with fresh media and ASO was added using a 3-point concentration response at 20uM, 4uM and 0.0274 uM. Cells were incubated with ASO for 7 days under gymnotic (free uptake) conditions.

qPCR was run per 384-well plate to calculate percent HTT remaining vs. mock-treated samples. Two biological replicates, each with two technical replicates, were run for each test article, with the biological replicates treated in separate 96-well plates. Percent HTT remaining vs. mock-treated samples parameter estimates were calculated using a robust linear mixed effects model controlling for the random intercept effects of technical replicate nested within biological replicate (robustlmm R package). Estimates of knockdown and standard error were derived from the model on a log2 scale and converted to a linear scale. For mock-treated samples, separate parameter estimates were derived for each mock well in the original 96-well compound plate, and a precision-weighted mock mean and standard error were calculated across all wells per experiment. The NTC was run across multiple experiments (N=8), so a precision-weighted NTC mean and standard error were calculated across all experiments.

Primary forebrain mouse neurons from Q111 mice were cultured as described (*75*). Briefly, forebrains were harvested from E18 mouse pups plated at a density of 250,000 cells/well onto 6-well poly-d-lysine coated dishes and fed glial conditioned media with B27 supplement (Thermo Fisher). Cells were incubated with ASO on post-plating days 2-10 before harvesting for downstream analysis.

### HTT protein isoform quantification

For MSD assay, lysates were prepared in a non-denaturing lysate buffer as previously described (*76*). For HTRF assay, a 10% (w/v) total protein homogenate was prepared and processed exactly as described (*30, 48*). Lysate dilutions and antibody concentrations are summarized in Supplementary Table S1. Western blots for HTT (Fig. 1) were run as described (*77*) and probed with anti-HTT antibody EPR5526 (Millipore; 1,2000). IPs (Fig. 3) were run as described (*49*) by IP with anti-polyQ antibody 3B5H10 (Millipore) and ID by anti-HTT antibody D7F7 (Cell signalling technologies).

### Immunohistochemistry

Brains from mice were collected and immediately post-fixed in formalin at 4°C for 18-20 hours before being paraffin embedded, sectioned at 5 um thickness, and mounted on super-frost glass slides. To perform immunostaining, sections were deparaffinized and rehydrated to distilled water as described previously(*76*). Primary antibodies: mouse anti-HTT1a (MW8; DSHB; 1:750), rabbit anti-NeuN (1:750; Millipore; ABN78) sheep-anti-HTTex1 (S830, 1:2000). Secondary antibodies: Alexa 488 anti-mouse (1:1000; Life Technologies), Alexa 568 anti-rabbit (1:1000; Life Technologies), Alexa 488 anti-sheep (1:500; Life Technologies). For MW8 images, 12-bit images were acquired with an IX-81 laser-scanning confocal microscope with Fluoview 1000 software (Olympus) using a 40x/1.30 NA oil objective. For S830 images, 8-bit images were acquired with a Zeiss LSM 980 confocal microscope with Zen blue software (Zeiss) using a 20x/0.8 NA objective.

For each secondary antibody, acquisition parameters were set such that a brain section with no primary antibody emitted no fluorescent signal and settings were maintained for the entire set of sections. Z-stack numbers and thickness were kept consistent for each set of sections. Maximum z-projections were compiled using ImageJ (*78*). NeuN and/or DAPI masks were created using ImageJ with custom macros and used to measure neuronal and/or nuclear expression of MW8 or S830.

### RNA sequencing

cDNA libraries were constructed at Azenta (South Plainfield, NJ) using the Illumina TruSeq RNA Sample Prep Kit with ERCC spike-in and sequenced on a HiSeq 2000 (2 x 150bp) to an average read depth of 2.9e7. Sequence reads were trimmed to remove adapter sequences and nucleotides with poor quality using Trimmomatic v.0.36. Reads were aligned to the *Mus musculus* GRCm38 ERCC reference genome assembly (ENSEMBL) using STAR aligner v.2.5.2b. Unique gene hit counts were calculated using featureCounts from the Subread package v.1.5.2. Differential gene expression analysis was conducted in R using edgeR (*79*), with voom (*80*). Unsupervised hierarchical clustering was performed using R package ‘pheatmaps’. The GO enrichment plots from Fig 7C were generated with package ‘cluterProfiler’ (*81*).

### Statistics

Experiments with multiple levels and groups were analyzed using factorial ANOVAs and post-hoc analyses were completed with Tukey HSD. All data and R scripts for analysis are available at the dryad repository https://doi.org/10.5061/dryad.6wwpzgn8j. RNAseq files are available at GEO accession GSE296071.

## List of Supplementary Materials

Table S1. Antibody dilutions

Table S2. Interventional experimental cohort details

Figure S1. MutASO induces elevation of NEFL

Figure S2. No behavioral differences between saline treated WT and Q111 mice

Figure S3. MutASO induces mild immune transcriptional signature in WT mice

## Acknowledgments

The authors thank Velvet Smith and Sage Berry for assistance with animal studies (Western Washington University), Andrea Grindeland, Deborah Cabin, and Jill O’Moore for their help with early pilot studies (McLaughlin Research Institute), David Howland, Thomas Vogt, and Deanna Marchionini for helpful comments (CHDI Foundation). Wave Life Sciences designed, synthesized, formulated and provided all oligonucleotides used in this work. They also provided the data and information shown in Figure panels 1B and 1D.

## Author contributions

Conceptualization: JBC

Methodology: RMB, CL

Investigation: RMB, CL, GO, EJS

Formal analysis: RMB, CL, GO

Funding acquisition: JBC, GPB

Supervision: JBC, GPB, CL

Writing – original draft: JBC, RMB

Writing – review & editing: JBC, RMB, CL, GPB

## Data and materials availability

Wave Life Sciences provided all ASOs used in the analysis and reasonable requests for these research materials for the purpose of directly replicating this procedure may be made by requesting a pertinent Material Transfer Agreement (MTA).

