## Supplemental material for "Selective targeting of mutant *huntingtin* intron-1 improves rescue provided by antisense oligonucleotides"

Table S1. Antibody dilutions

| **Assay** | **Protein species** | **Antibody** | **Acceptor** | **Dilution** |
|---|---|---|---|---|
| HTRF | Soluble mHTT (excluding HTTexon1) | MAB2166-Tb : 4C9-488 | 20 ng | 5% (1 in 2) |
| HTRF | Soluble HTTexon1 | 2B7-Tb : 1B12-d2 | 40 ng | 10% |
| HTRF | HTTexon1 aggregation | 4C9-Tb : 1B12-d2 | 40 ng | 10% |
| MSD | mHTT | 2133 : D7F7-ST | N/A | N/A |
| MSD | wtHTT | 2B7 : MW1-ST | N/A | N/A |
| Western Blot | HTT | EPR5526 | N/A | N/A |
| IP | HTT IP | 3B5H510 | N/A | N/A |
| IP | HTT IP-detection | D7F7 | N/A | N/A |

Table S2. Interventional cohort mouse details

| Genotype | Treatment | Sex | N | Endpoint |
|---|---|---|---|---|
| WT | Saline | F | 7 | 7 |
| WT | Saline | M | 8 | 8 |
| WT | PanASO | F | 7 | 6 |
| WT | PanASO | M | 8 | 5 |
| WT | MutASO | F | 8 | 8 |
| WT | MutASO | M | 7 | 6 |
| Htt.Q111/+ | Saline | F | 8 | 8 |
| Htt.Q111/+ | Saline | M | 8 | 8 |
| Htt.Q111/+ | PanASO | F | 7 | 7 |
| Htt.Q111/+ | PanASO | M | 8 | 8 |
| Htt.Q111/+ | MutASO | F | 8 | 8 |
| Htt.Q111/+ | MutASO | M | 7 | 7 |
| | | Total | 91 | 86 |

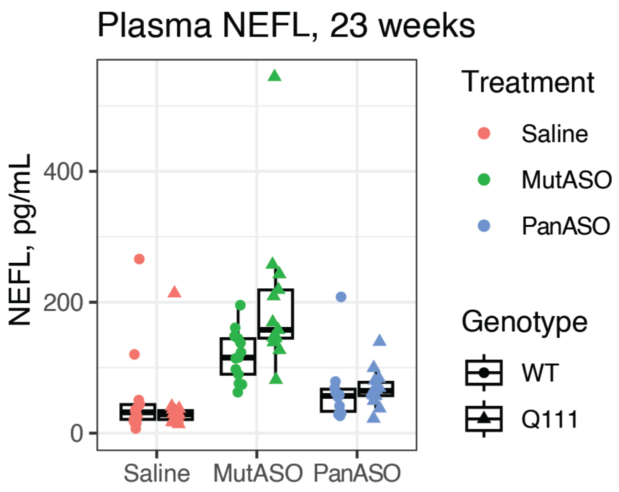

**Figure S1. MutASO induces elevation of NEFL.** MutASO induced a 3.5 fold increase in NEFL levels compared to saline (Mean MutASO NEFL = 158.5 pg/mL; Mean Saline NEFL = 45.8; ANOVA $p < 0.0001$; Tukey $p < 0.001$), with elevation higher in Q111 than WT (Mean WT MutASO = 118 pg/mL; Mean Q111 MutASO = 199 pg/mL; ANOVA interaction $p = 0.02$). PanASO did not alter NEFL levels compared to saline (ANOVA $p = 0.44$).

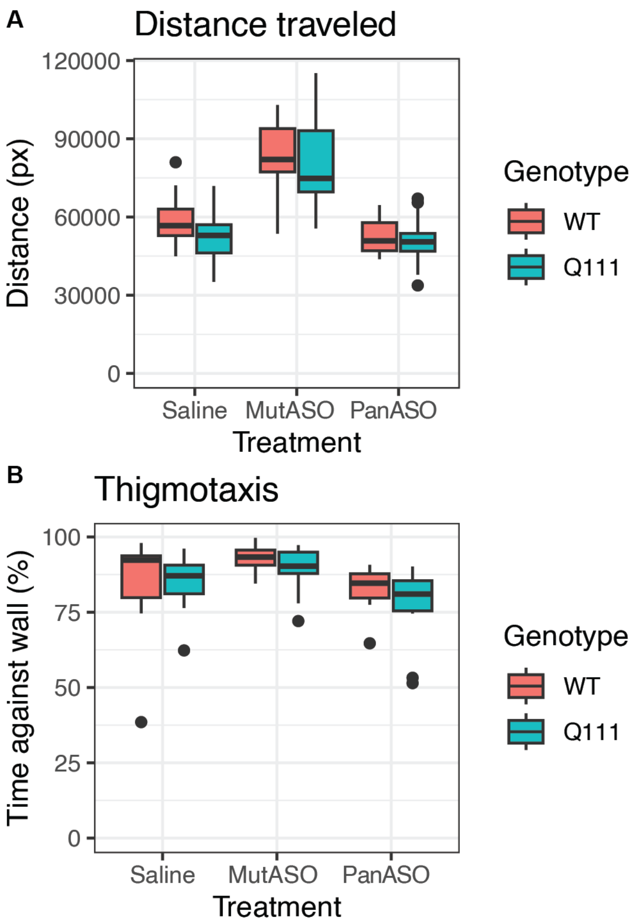

**Figure S2. No behavioral differences between saline treated WT and Q111 mice.** ANOVA for both revealed significant effect of genotype and treatment ($p < 0.05$) on distance traveled, however post-hoc analyses revealed (A) distance traveled (Tukey HSD: Q111 saline vs WT saline, $p = 0.87$, n/group = 15-16) and (B) thigmotaxis (Tukey HSD: Q111 saline vs WT saline, $p = 0.99$, n/group = 15-16) were not altered at baseline.

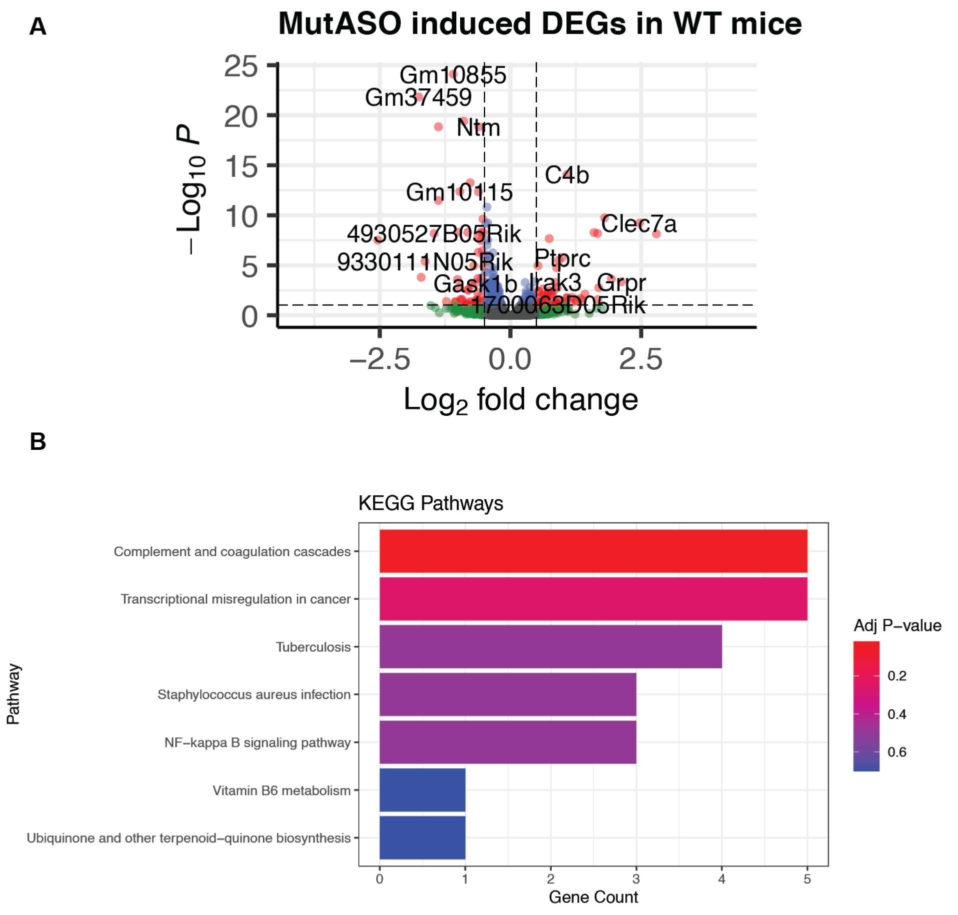

**Figure S3. MutASO induces mild immune transcriptional signature in WT mice.** (A). Volcano plot of 119 DEGs (FDR < 0.1, absolute Log2FC > 0.5) induced by MutASO in WT mice. (B) KEGG pathway enrichment reveals one significantly enriched pathway (adjusted p = 0.02) with 5 genes. Additional pathways that did not meet significant (p > 0.05) adjusted threshold are shown.
